# The structure of immature tick-borne encephalitis virus

**DOI:** 10.1101/2023.08.04.551633

**Authors:** Maria Anastasina, Tibor Füzik, Aušra Domanska, Lauri IA Pulkkinen, Lenka Šmerdová, Petra Pokorná Formanová, Petra Straková, Jiří Nováček, Daniel Růžek, Pavel Plevka, Sarah J Butcher

## Abstract

Tick-borne encephalitis virus (TBEV) is a medically important flavivirus that poses a significant health threat in Europe and Asia. However, the structure of the immature form of TBEV remains unknown. Here, we employed state-of-the-art cryogenic electron microscopy (cryoEM) to determine the structure of the immature TBEV particle. The immature TBEV particle has a diameter of 56 nm and its surface glycoproteins are organised into spikes characteristic of immature flaviviruses. The cryoEM reconstructions of the whole virus and of the individual spike enabled us to build atomic models of the major viral components, the E and prM proteins. The insights obtained from our study provide a foundation for understanding the early stages of TBEV assembly and maturation. The pr domains of prM have a critical role in holding the heterohexameric prM3E3 spikes in metastable conformation. Destabilisation of the prM furin-sensitive loop at acidic pH facilitates its processing. The prM cleavage, the collapse of E protein ectodomains onto the virion surface concurrent with significant movement of the membrane domains of both E and M, and release of the pr fragment from the particle render the virus mature and infectious. This knowledge contributes to our understanding of the flavivirus life cycle.

## Introduction

Tick-borne encephalitis virus (TBEV; *Orthoflavivirus encephalitidis*) belongs to the genus *Orthoflavivirus* and infects a range of ticks, birds, and mammals, including humans. The symptomatic infection in humans results in tick-borne encephalitis (TBE), a severe neurological disease that often results in long-term sequelae and can be fatal (reviewed in ^1^). The disease manifestations vary depending on the virus subtype, with the European subtype causing milder disease with a 0.5–2% mortality rate, the Siberian subtype often causing long-term or chronic infections with 1–3% fatality rate, and the Far Eastern subtype having the highest death rate of up to 35 %^2^. TBEV is endemic within Europe, Russia, and North Eastern Asia^3^. Despite the available vaccines, the number of infections has been steadily growing over the last few decades with about 10,000–15,000 annual reported cases (reviewed in ^4^). Specific therapies for TBE are not available.

TBEV is structurally similar to other flaviviruses^5,6^, but has been less well studied than its mosquito-borne counterparts such as dengue (DENV), Zika (ZIKV), West Nile (WNV), Japanese encephalitis (JEV) and yellow fever (YFV) viruses. The virion consists of a single-stranded positive-sense RNA genome bound to multiple copies of the capsid (C) protein, surrounded by a host-derived lipid bilayer where 180 copies of membrane (M) and envelope (E) proteins are embedded forming an icosahedrally-symmetric shell. The building block of the shell in a mature virus is an E_2_M_2_ heterotetramer, with 90 such units lying parallel to the membrane in a smooth “herringbone” arrangement^5,6^. TBEV enters the cells via receptor-mediated endocytosis where E plays a major role, mediating receptor binding and low pH-induced membrane fusion^7,8^. Following uncoating, the viral genome is translated into a single multipass membrane polyprotein that is proteolytically processed to yield individual structural and non-structural proteins. Viral RNA-dependent-RNA-polymerase synthesizes negative-sense and subsequently positive-sense copies of the genome^9^.

The newly synthesised genomes bind to multiple copies of C to form nucleocapsids, which acquire an envelope by budding into the lumen of the endoplasmic reticulum. The envelope of immature TBEV particle contains 180 copies of each M protein precursor (prM) and E forming heterodimers. The prM protein stabilises E and covers its fusion loop, preventing premature virus activation and its fusion in the low pH with the membranes of the *trans*-Golgi network (TGN)^10,11^. Immature particles of most flaviviruses are fusion-incompetent and non-infectious and must mature to acquire infectivity^7,12^. Flavivirus maturation occurs at mildly acidic pH, when host protease furin cleaves the prM protein to a pr peptide that will later dissociate from the particle, and the M protein that remains in the particle, resulting in infectious viruses ^13,14^.

Particle heterogeneity^15^, flexibility^16^, and symmetry imperfections^17^ have limited the achievable resolution of immature flavivirus particle reconstructions. Detailed structural information is available only for the mosquito-borne Spondweni (SPOV) and mosquito-specific Binjari (BinJV) viruses, providing the first atomic details about the prME interactions^18,19^.

Here, we used cryogenic electron microscopy (cryoEM), single particle analysis, and localised reconstruction to determine the structure of immature TBEV particles^20–24^. By analysing two model TBEV strains—Hypr and Neudoerfl—and the Kuutsalo-14 isolate, we obtain a comprehensive information about the architecture and molecular organisation of immature TBEV, and propose the major conformational changes that need to occur on the route to maturation.

## Results and Discussion

To acquire a comprehensive understanding of the immature TBEV particle structure, we investigated three strains: Hypr, Neudoerfl, and Kuutsalo-14, using independent protocols for particle production, cryoEM data collection, and image processing. The immature particles were produced in infected cells treated with ammonium chloride to increase the pH of the exocytic pathway and thereby inhibit maturation^12,13,25^. The purified samples contained primarily immature particles as indicated by the presence of a pronounced prM band on SDS-PAGE (Fig. S1). Prior to vitrification for cryoEM, we inactivated the virus using formaldehyde or ultraviolet (UV) light that neutralised virus infectivity without affecting protein structure^6,26^.

The cryoEM micrographs of all three strains revealed particles with a spiky appearance characteristic of immature flaviviruses (Fig. 1a). However, we observed considerable heterogeneity in particle structure, including broken particles, and particles with mixed spiky and smooth morphologies, which may represent partially mature particles. Reference-free 2D class averages revealed protein densities projecting outward from the particle surface, causing the spiky morphology. However, in some of the 2D classes, one side of the particle was blurred, with poorly resolved protein and lipid bilayer densities. This blurring could be attributed to sample heterogeneity and/or symmetry imperfections within the particles (Fig. 1a). Imposition of icosahedral symmetry during the reconstruction enabled calculation of 7.1 Å-resolution maps of Kuutsalo-14 and Neudoerfl, and an 8.6 Å-resolution map of Hypr according to the 0.143 gold-standard criterion of Fourier shell correlation (GSFSC) cutoff^27^ (Fig. 1b and S2-S4, Table 1). The reconstructed particles contain 60, nonsymmetrical, prM_3_E_3_ “spikes” (Fig. 1b).

**Figure 1.**
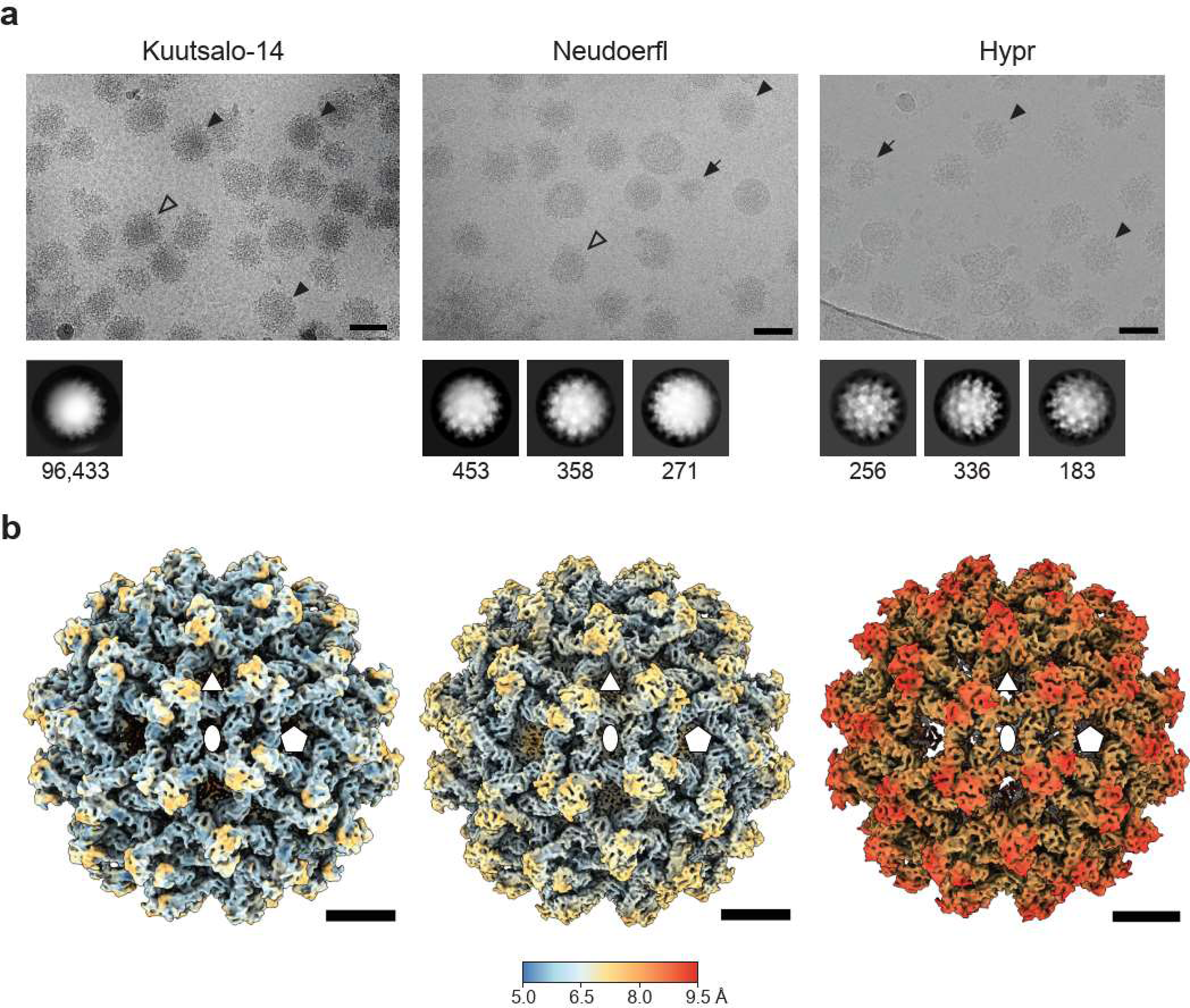
CryoEM and 3D reconstructions of immature TBEV particles. **(a)** representative micrographs, where black arrowheads indicate spherical particles, white arrowheads indicate non-spherical particles and black arrows indicate broken particles. Scale bar 50 nm. 2D class averages box sizes are 800 x 800 Å for Kuutsalo-14 and 864 x864 Å for Neuroerfl and Hypr. The prevalent initial 2D class averages are shown below the respective micrographs with the number of particles per class indicated. (**b**) Isosurface representations of the icosahedral-symmetrised 3D reconstructions of immature Kuutsalo14, Neudoerfl, and Hypr particles viewed down an icosahedral twofold axis of symmetry,coloured by local resolution with the key indicated by the bar. Positions of selected symmetry axes are indicated by a pentagon (fivefold), an ellipse (twofold), and a triangle (threefold). Scale bar 10 nm.

**Table 1.** Cryo-EM data collection, refinement, and validation statistics.

|  | Kuutsalo-14<br>full particle | Kuutsalo-14<br>prM<sub>3</sub>E<sub>3</sub> spike | Neudoerfl<br>full particle | Neudoerfl<br>prM<sub>3</sub>E<sub>3</sub> spike | Hypr<br>full particle |
| --- | --- | --- | --- | --- | --- |
| **Data collection and processing** |  |  |  |  |  |
| Magnification | 81000 | 81000 | 105000 | 105000 | 75000 |
| Voltage (kV) | 300 | 300 | 300 | 300 | 300 |
| Electron exposure (e<sup>-</sup>/Å<sup>2</sup>) total dose | 43 | 43 | 40 | 40 | 69 |
| Defocus range settings (μm) | -0.7 to -2.7 at 0.2 step | -0.7 to -2.7 at 0.2 step | -1.0 to -3.0 at 0.2 step | -1.0 to -3.0 at 0.2 step | -1.0 to -3.0 at 0.2 step |
| Pixel size (Å) | 1.075 | 1.075 | 1.563 (binned 1.875x) | 0.8336 | 1.62 (binned 1.5x) |
| Symmetry imposed | I2 | C1 | I1 | C1 | I1 |
| Micrographs (no.) | 40138 | 40138 | 11246 | 11246 | 2262 |
| Initial particle images (no.) | 271870 | 4312440 | 156614 | 1639578 | 29978 |
| Particles used in reconstruction (no.) | 70189 | 1172996 | 36236 | 552993 | 18160 |
| Map resolution (Å) | 7.1 | 3.9 | 7.0 | 4.0 | 8.6 |
| FSC threshold | 0.143 | 0.143 | 0.143 | 0.143 | 0.143 |
| Map resolution range (Å) | 999-7.1 | 999-3.9 | 999-7.2 | 999-4.0 | 999-8.6 |
| **Refinement** |  |  |  |  |  |
| Map sharpening B factor (Å<sup>2</sup>) | -658.6 | -226.9/variable | -725 | -167 | -1178 |
| **Model composition** |  |  |  |  |  |
| Non-hydrogen atoms | N/A | 14700 | N/A | 13407 | N/A |
| Protein residues | N/A | 1923 | N/A | 1908 | N/A |
| E | N/A | 1-496 | N/A | 1-492 | N/A |
| prM | N/A | 1-82, 85-98, 112-162 | N/A | 1-82, 86-95, 111-160 | N/A |
| Ligands | N/A | 3 (NAG) | N/A | 3 (NAG) | N/A |
| **R.m.s. deviations** |  |  |  |  |  |
| Bond lengths (Å) | N/A | 0.66 | N/A | 0.002 | N/A |
| Bond angles (°) | N/A | 1.06 | N/A | 0.455 | N/A |
| **Validation** |  |  |  |  |  |
| MolProbity score | N/A | 1.27 | N/A | 1.57 | N/A |
| Clashscore | N/A | 1.57 | N/A | 5.31 | N/A |
| Poor rotamers (%) | N/A | 0.06 | N/A | 0.00 | N/A |
| **Ramachandran plot** |  |  |  |  |  |
| Favored (%) | N/A | 94.68 | N/A | 95.85 | N/A |
| Allowed (%) | N/A | 5.32 | N/A | 3.99 | N/A |
| Disallowed (%) | N/A | 0.00 | N/A | 0.16 | N/A |
| PDB ID | N/A | 8PPQ | N/A | 8PUV | N/A |
| EMD | 17809 | 17808 | 17946 | 17947 | 17945 |

The structure of the nucleocapsid is not resolved in the cryoEM reconstructions of immature TBEV particles indicating that it does not follow icosahedral symmetry. However, the transmembrane helices of E and prM traversing the lipid bilayer can be readily observed in cross-sections of the icosahedrally-symmetric reconstructions (Fig. 2a-c). The ectodomains of E and prM form the spikes extending outwards from the particle surface, contributing to the larger diameter of immature TBEV compared to the mature virion (∼560 Å vs. ∼500 Å; Fig. 1b, 2a-d). The mature virion exhibits a distinctly angular inner membrane leaflet, due to the clustering of the E and M protein transmembrane helices (Fig. 2d). In contrast, both membrane leaflets in the immature particle are round and the inner leaflet is more closely juxtaposed to the NC. This striking difference in the membrane shape is due to the different spatial organisation of the transmembrane domains in mature and immature TBEV (Fig. 2e-f). Furthermore, the transmembrane helices of E and M are slanted within the membrane in immature particles (Fig. 2c, 3b), contrasting to their orthogonal insertion in the virion membrane (Fig 2d). Evidently, the topology of E differs between immature and mature particles, manifesting in unique arrangements of the ectodomain and membrane-associated regions, and greater projection of the E ectodomains outwards from the particle surface in the immature TBEV. The membrane helix domains of both E and M can be unambiguously connected to their ectodomains in all three reconstructions. The connections in prM are similar to those found in BinJV (Fig. S5), but not to those presumed in the immature Spondweni, Zika or DENV particles^18,19,28,29^.

**Figure 2.**
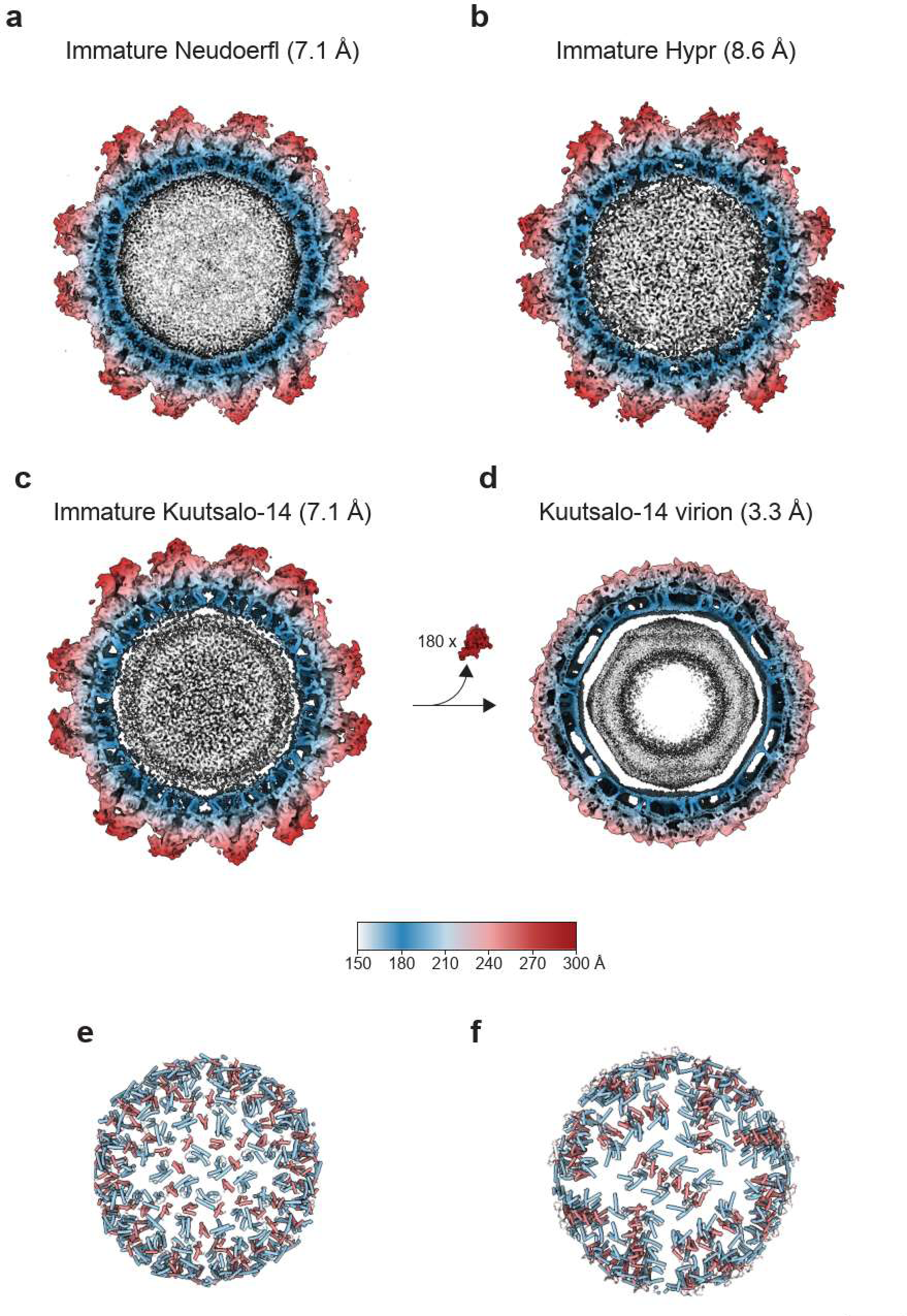
Membrane organisation in immature vs mature TBEV particles. (**a-c**) 100 Å thick central sections of icosahedral reconstructions of immature particles compared to (**d**) the mature Kuutsalo-14 reconstruction (EMDB ID:14512)^6^. Cleavage and dissociation of 180 copies of pr peptide upon maturation is indicated between **c** and **d**. (**e**) Positions of transmembrane and peripheral membrane helices of E (light blue) and (pr)M (light red) in the front hemisphere of the immature Kuutsalo-14 and (**f**) in the mature Kuutsalo-14. Membrane-associated helices cluster into rafts in the mature particle compared to the more even distribution in the immature particle. The scale bar is 10 nm.

The interpretability of the icosahedral reconstructions of the whole immature TBEV particles was limited by the resolution (Fig. 2a-c, 3a, S4, Table 1). However, one can clearly identify the three prME heterodimers forming individual spikes at the particle surface (Fig 1b, 2a-c, 3a). Suspecting the resolution was limited by particle heterogeneity (Fig. 1a), we employed localised reconstruction of individual spikes, and were able to improve the resolution for Kuutsalo-14 to 3.9 Å, and the resolution of Neudoerfl to 4.0 Å at the 0.143 FSC cutoff (Fig. S4), and to build atomic models for both strains. We observed an 1.0 Å root-mean-square deviation (RMSD) between Kuutsalo-14 and Neudoerfl atomic models when the Cα of prM and E ectodomains were compared and 2.1 Å RMSD of the Cα of the membrane-associated helices emphasising the reliability of the independently determined structures (Fig. S6). Localised reconstruction of Hypr did not result in an improved resolution compared to the icosahedral reconstruction and was not pursued further.

Each spike decorating the surface of immature TBEV particle is formed by three prME heterodimers and is asymmetric with two prME opposing each other (Fig. 3b-c, red and yellow dimers), and a third dimer joining from a side (Fig. 3b-c, blue dimer; Supplemental video). The peripheral (PM) and transmembrane (TM) helices of prM and E embed the proteins in the lipid bilayer. The E ectodomains extend outwards from the particle surface joining at the top like a tripod, with the pr domains of prM positioned at the top (Fig. 3 a,b). The ectodomains of pr and E in the immature TBEV display the characteristic fold of flavivirus proteins, where pr comprises a globular beta-sandwich and E consists of three distinguishable domains (DI, DII, DIII) (Fig. 3b, S7). Our models reveal a discrepancy in the E ectodomain topology compared to the recently published prME crystal structure^11^ (Fig. S7). In the latter, E-DII adopts a different position relative to DI resulting in an 19.5° difference with a total Cα RMSD of 8.6 Å between the models^11^. The difference may arise due to the different oligomeric states of purified E ectodomains compared to the native E proteins within the particle, or due to the different probed pH. These differences underline the importance of structural data obtained within the whole virus particle context, both at neutral and acidic pH.

**Figure 3.**
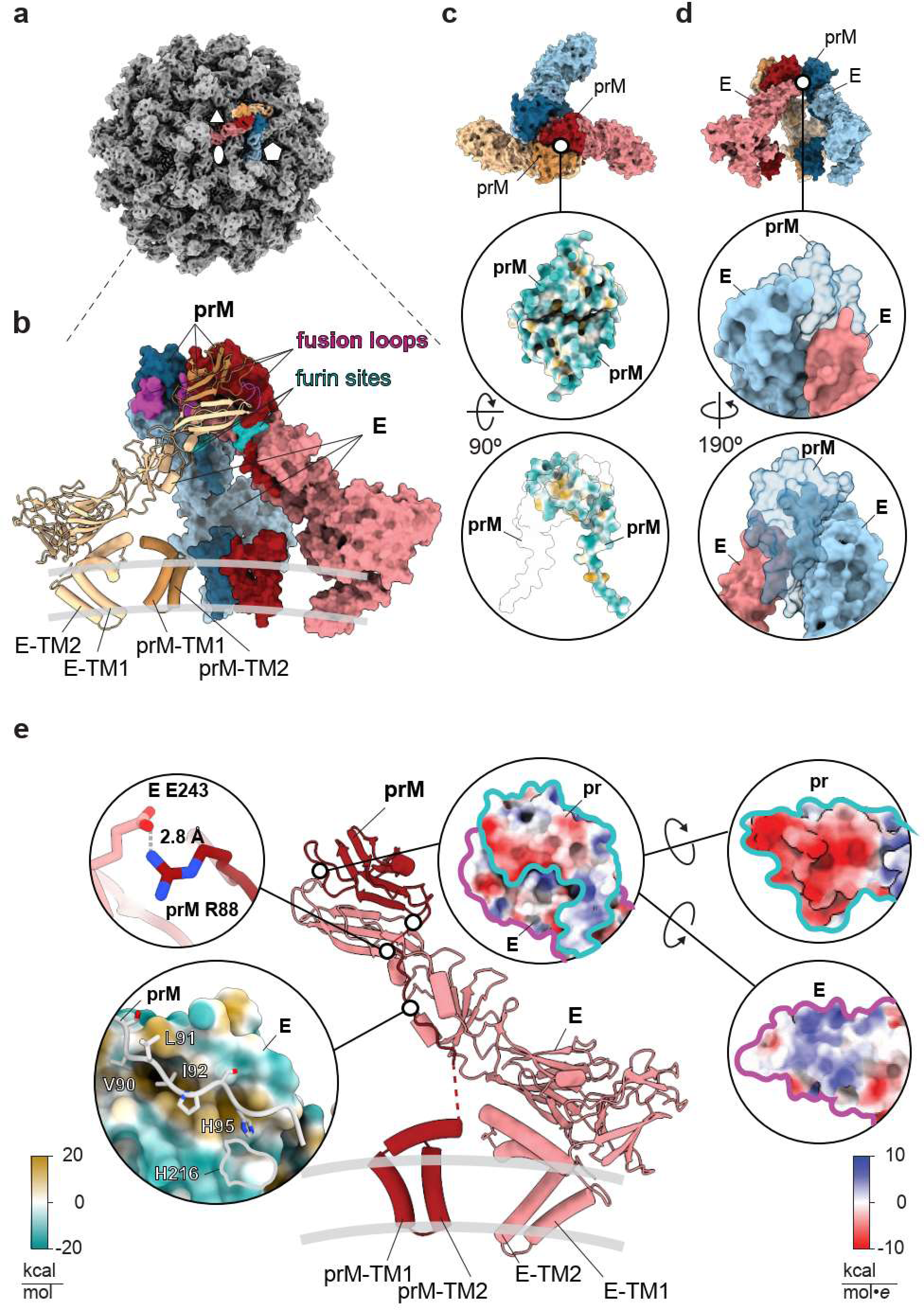
prM_3_E_3_ spike organisation and protein-protein interactions. (**a**) Isosurface representation of the icosahedral reconstruction of immature Kuutsalo-14. Positions of symmetry axes are indicated using an oval (twofold), a triangle (threefold), and a pentagon (fivefold). One prM_3_E_3_ spike is highlighted in colour. (**b**) Atomic model of a Kuutsalo-14 prM_3_E_3_ spike refined against a 3.9 Å resolution map. One prME heterodimer is shown in cartoon representation with TM helices of E and prM indicated. The remaining two prME heterodimers are shown as molecular surfaces. prM and E dimers are coloured in red, yellow, and blue with prM in darker shades. Fusion loops of E and prM furin cleavage sites are highlighted in magenta and turquoise, respectively. (**c**) A molecular surface representation of prM_3_E_3_ with a close-up view of the pr-pr interaction interface. Proteins are coloured as in (**b**) in the top panel, both pr peptides are coloured by hydrophilicity in the middle, and in the bottom panel one pr peptide is coloured according to hydrophilicity whereas the other prM is shown as a transparent molecular surface with black outline to indicate the interaction area. (**d**) A molecular surface representation of prM_3_E_3_ spike with a close-up view of E-prM-E interaction interfaces showing how one prM binds together the two E. The proteins are coloured as in (**b**), but prM is semi-transparent. (**e**) A cartoon representation of one prME heterodimer with names of TM helices of pr and E indicated. Close-up images show prME salt bridge (left upper panel), a hydrophobic zipper (left, bottom) stabilising the furin site, and prME interaction interface coloured by electrostatic potential (right side; proteins are outlined for clarity).

The E-DII harbouring the fusion loops join at the top of the prM_3_E_3_ spike and are capped by pr peptides, each of which binds to its respective E through an interface with a cumulative buried area of ∼4700 Å^2^. The negatively-charged prM surfaces cover the positively charged E-DII tips, obscuring the fusion loops (Fig. 3b,e, Supplemental video). The heterohexamer structure of the spike is stabilised by pr-pr interactions, as direct E-E contacts in the spike are limited to just one small interaction surface of ∼160 Å^2^ (Supplemental video). The two opposing prME dimers interact via a symmetrical pr-pr interface with ∼460 Å^2^ buried surface area (Fig. 3c), involving hydrophobic residues Ala19, Ala20, Val31, Leu33, Val59, Val61, and Phe64 of both prM proteins. The third prME dimer joins from a side via a ∼160 Å^2^ interface between its E protein and E from another dimer, and ∼320 Å^2^ interface between its pr and E of another dimer (Fig. 3d). Hydrophobic residues are enriched in both interfaces. Thus, the prM_3_E_3_ arrangement of the immature TBEV particle is mostly maintained by the interactions between pr domains within the prM_3_E_3_ spike. Analysis of E and M protein sequences of 182 TBEV isolates belonging to all three virus subtypes indicates that the interacting residues are highly conserved^6^.

The globular pr domain of prM is connected to the membrane-associated helices by a flexible linker (Fig. S5). We were able to build the linker along the E protein up to Gly98, including the conserved 85-Arg Thr Arg Arg-88 furin recognition site^14^. The cleavage of prM at Arg88 splits it into pr peptide (residues 1-88) and M protein (residues 89-162) and is a crucial step of TBEV maturation^14^. The residues forming the furin cleavage site are held in place by a polar interaction between Arg88 of pr and Glu243 of E, and is zipped downstream by the hydrophobic residues Val90, Leu91, and Ile92 of prM fitted into a hydrophobic pocket on E (Fig. 3e). This hydrophobic zipper is maintained in mature TBEV, where Val2, Leu3, and Ile4 of M (corresponding to Val90, Leu91, and Ile92 of prM) are docked into a hydrophobic pocket of E underneath E-DII (Fig. S8), suggesting that the interaction is stable throughout the conformational rearrangement of prME and prM processing. The location of the M protein N-terminus beneath the E-DII suggests that prM cleavage precedes the formation of the mature particle’s herringbone structure, as the furin cleavage site would otherwise be inaccessible. Comparing to the low pH structure of a fragment of pr complexed with preassembled E ectodomain dimers^11^, even there, the furin cleavage site would be inaccessible from the surface of the virus.

Within the immature particle context, the furin cleavage sites are located “inside” the spike (Fig. 3B, Supplemental video) and are inaccessible to the globular 88 kDa furin^30^ without a conformational rearrangement of the spike. *In vitro*, the susceptibility of prM in immature TBEV to furin cleavage increases at pH 7 and below, indicated by a reduction of the prM, and the emergence of the M protein bands on SDS-PAGE, verified by Western Blot (Fig. 4). The proteolysis is likely facilitated by the conformational changes of the spike that increase the cleavage site accessibility, as furin is known to be efficient throughout all our tested pH values^13,31^ (Fig 4). This pH-triggered conformational change of the spike agrees with earlier observations of pH-induced changes in the immature particle antigenicity^10^. There are 15 conserved His in TBEV prME. A strictly conserved His95 from the prM linker interacts with the E hydrophobic pocket (Fig. 3e). This histidine is located in proximity to another conserved histidine, His216 in E, and we speculate that when protonated, these residues could trigger exposure of the loop containing the furin cleavage site to allow proteolysis. Protonation of histidines in the prM linker has been proposed as a maturation trigger for DENV (His98), SPOV (His101), and BinJV (His88)^18,19,32^, however, the exact mechanism is yet to be shown.

**Figure 4.**
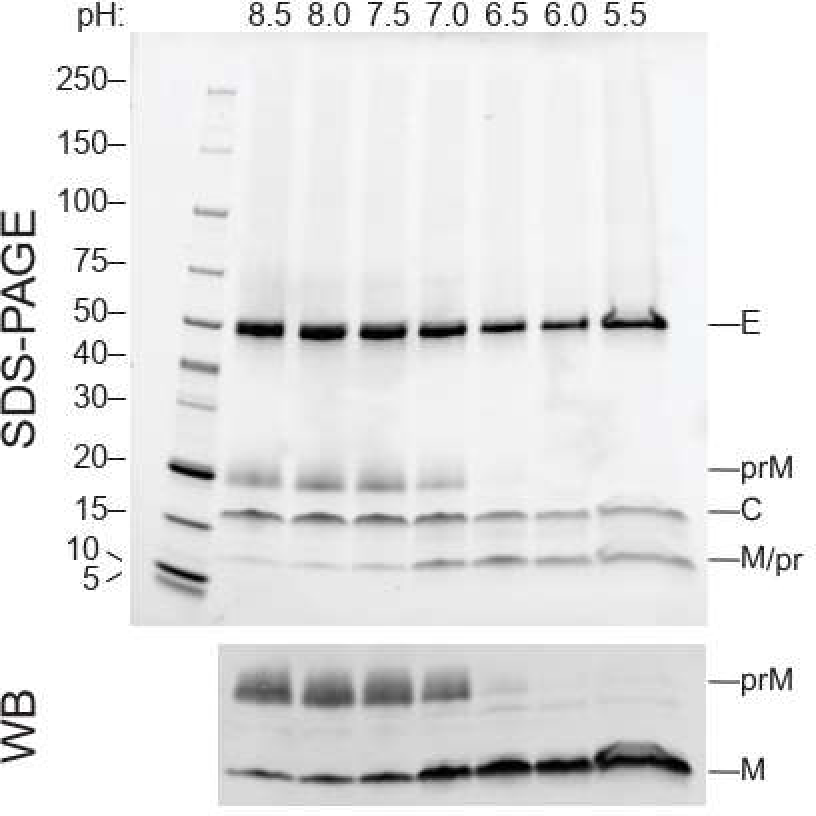
Susceptibility of immature TBEV to furin at decreasing pH. An SDS-PAGE of purified immature TBEV shows the three major protein components, E, prM and C. As the particles are exposed to decreasing pH in the presence of furin for 16 h at 30 °C, prM is cleaved into pr peptide and M with increasing efficiency (top panel), which is verified by immunostaining with an antibody against the C-terminus of M, as pr and M have similar molecular weights (bottom panel).

The prM linker (residues 99-111) is flexible and is not resolved in our prM_3_E_3_ maps (Fig. 3b, Supplemental video). However, densities connecting the globular pr domain to membrane helices of prM are evident in the icosahedral reconstructions of all the three immature TBEV particles (Fig. S5), allowing us to assign the positions of prM membrane domains beneath the spike tripod (Fig. 3b). Importantly, none of the spikes in the icosahedral reconstructions sit directly on the imposed symmetry axes, and thus all the linkers can be treated as independent observations. A similar domain topology of prM was reported earlier for BinJV^19^, whereas an alternative model was suggested for SPOV^18^, which proposes clustering or prM and E membrane-associated domains (Fig. 5a,b). The overall TM domain distribution across the virus particle is, however, similar for all the three viruses (Fig. 5c). We propose that the membrane domain topology is similar in all of the immature flaviviruses with the evidence coming from both TBEV and BinJV (Fig. 5d-f), and that the connection has been incorrectly assigned for other viruses, including SPOV^18,28,29^. The correct assignment affects the interpretation of movements required for particle maturation.

**Figure 5.**
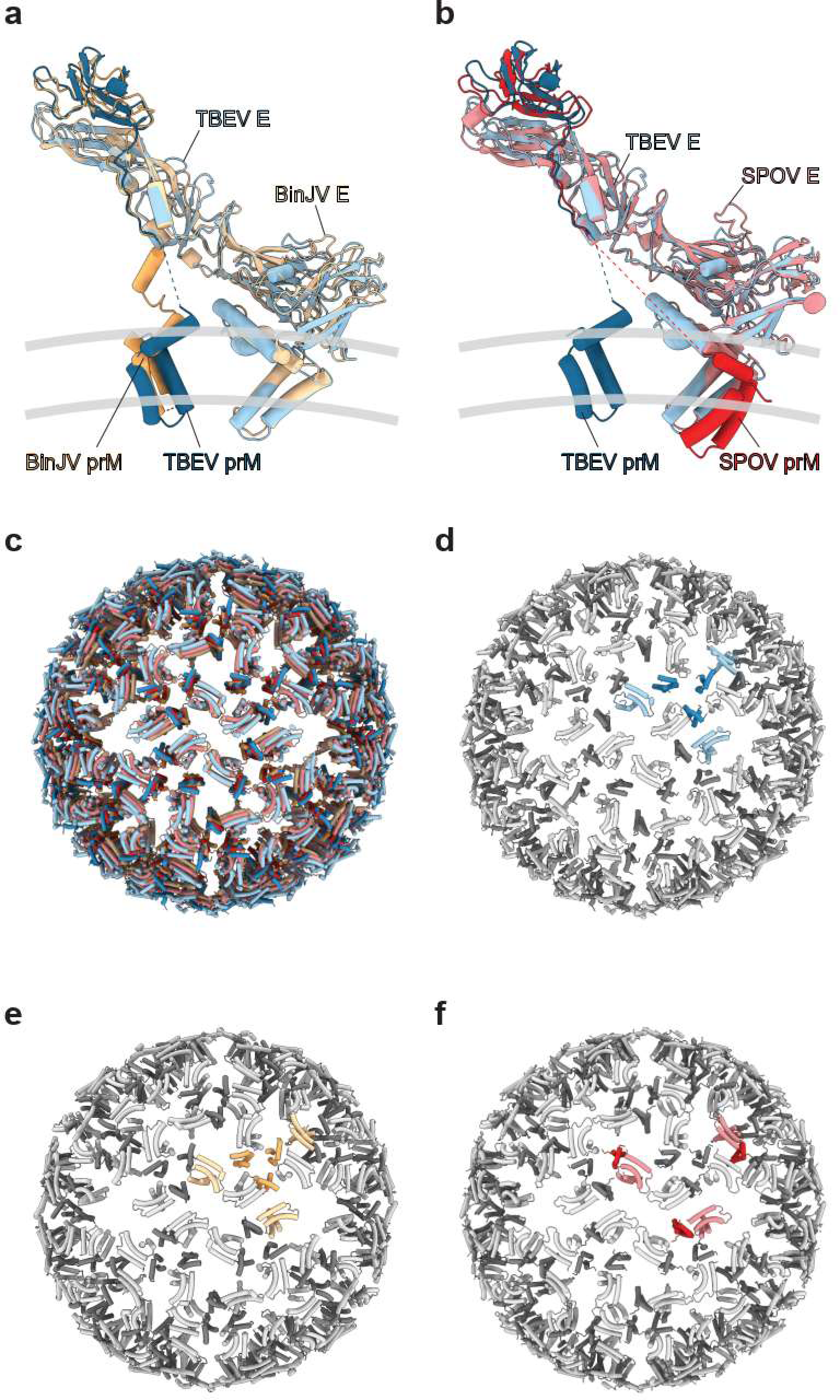
Comparison of membrane-associated domains of immature TBEV, BinJV and SPOV particles. Cartoon representations of Kuutsalo-14 prME (blue) overlaid with (**a**) BinJV (yellow; PDB ID: 7l30) or (**b**) SPOV (red; PDB ID: 6zqj) show that prM membrane-associated domains are localised underneath the spike in TBEV and BinJV, but are clustered with E membrane-associated domain in SPOV. (**c**) Only the transmembrane and peripheral membrane helices of E and prM are shown for the front hemispheres of immature TBEV (blue), BinJV (yellow) and SPOV (red) particles emphasising similar distribution of membrane-associated domains in all three viruses. (**d-f**) Only the transmembrane and peripheral membrane helices of E (tints) and prM (shades) are shown for the front hemispheres of immature TBEV (**d**), BinJV (**e**) and SPOV (**f**). Membrane-associated domains of E and prM of one asymmetric unit per particle are highlighted in colour, emphasizing similar domain assignment in TBEV (**d**, blue) and BinJV (**e**, yellow), but different in SPOV (**f**, red).

The mechanism of particle maturation is a central question in flavivirus biology, which may be addressed through the interpretation of immature and mature particle structures and aided by biochemical data^5,6,10,11,13,14,18,19,33^. The events driving maturation require destabilisation of prM-prM interactions at the tip of the spike, exposure of the furin site in prM and its cleavage, and collapse of E ectodomains onto the membrane accompanied by the large movement of the E and M TM domains. Our structural data support the collapse concept of flavivirus maturation, where E ectodomains reposition parallel to the particle membrane as soon as the spike is disrupted^19^. In TBEV, this conformational change is irreversible^13^. The immature particle heterogeneity (Fig. 2a), the prM_3_E_3_ spikes flexibility, and the pH-sensing histidines in the proximity of the furin cleavage sites (Fig. 3e) may facilitate local conformational changes at acidic pH to enable prM cleavage. As the ammonium chloride treatment of the cells used to produce immature particles raises the exocytic pathway pH, based on this and the *in vitro* furin sensitivity assay (Fig.4), the conformational changes enabling furin cleavage occur at pH 7 and below^13^.

After spike dissociation, the E dimer formation is probably driven by the large dimerisation surfaces of E proteins^33^. Within one spike, two E proteins (red and yellow in Fig. 3b,c and Fig. 6a,b) already have their dimerisation surfaces facing each other. Even though in the immature spike, these E proteins have no interactions with each other, when they collapse onto the membrane they would be readily positioned to dimerise (red and yellow E proteins in Fig. 6c). The third E of that spike (blue in Fig. 3b,c and Fig. 6a,b) will have to interact with the corresponding E of another spike, one such option is shown in Fig. 6a-c. So far there is no satisfactory molecular model of the immature to mature structure transition that we are aware of that does not involve clashing of the proteins, and has the appropriate assignment of all the transmembrane domains. Based on the large heterogeneity of the immature particles, we propose that rather than all spikes maturing at once, there is more likely to be nucleation centre of the conformational change from the most dynamic (disordered) region on the surface, even at pH7 and maturation propagates across the surface^34^. In so doing, there could be an additional opening up of the prME that allows better proteolytic access to the furin cleavage site than either of the two static models currently reveal. In conclusion, the prM cleavage, the collapse of E protein ectodomains onto the virion surface concurrent with significant movement of the membrane domains of both E and M, and release of the pr fragment from the particle render the virus mature and infectious. This knowledge contributes to our understanding of the flavivirus life cycle and can be aided with future studies on the pH-activated immature particle and timing of pr dissociation.

**Figure 6.**
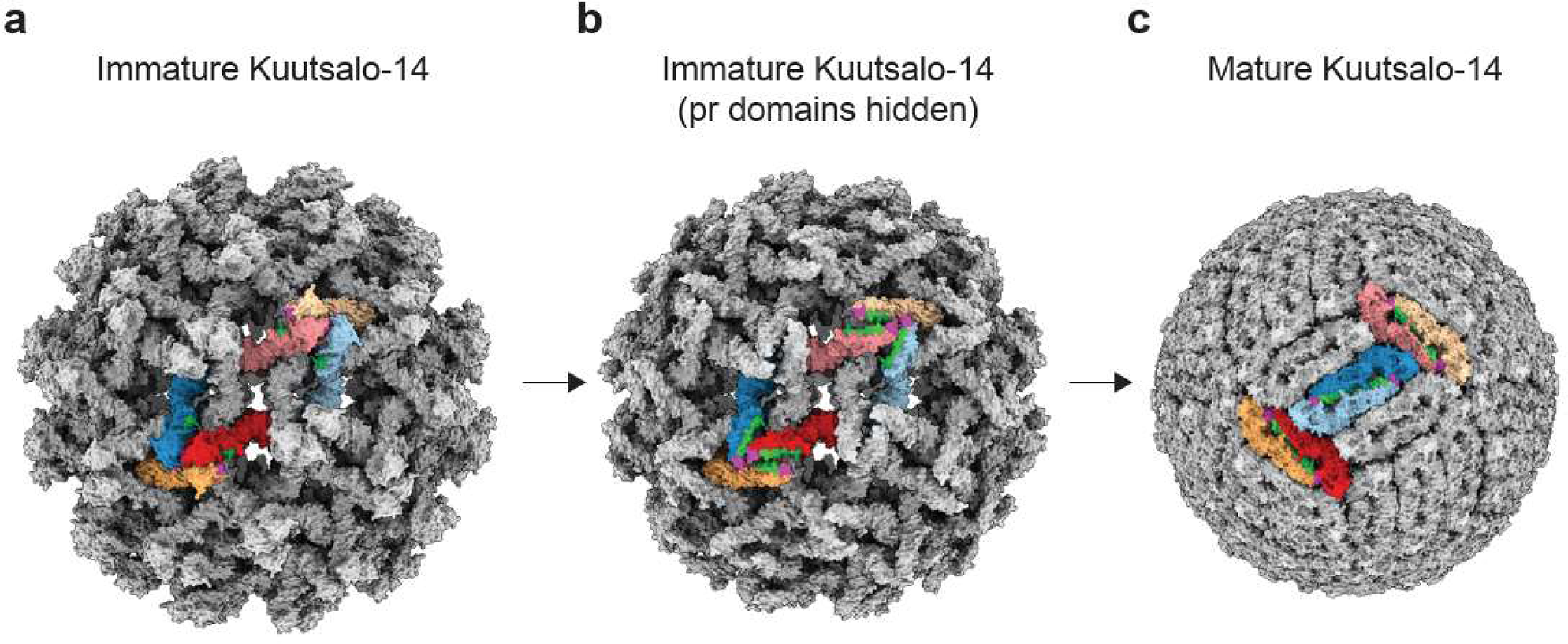
Redistribution of E ectodomains after TBEV maturation. (**a**) Surface representation of an immature Kuutsalo-14 particle is shown with two asymmetric units highlighted in colour, individual prME dimers are coloured in tints and shades of red, yellow and blue for clarity. (**b**) Same, but pr domains are hidden for clarity, and the residues of E that will form a E-E dimer interacting surface in the virion are highlighted in green, the fusion loops of E are highlighted in pink. (**c**) Proposed redistribution of E on the virion surface after maturation involves collapse and dimerization of two E from one prM_3_E_3_ spike (red and yellow), whereas the third E (blue) will interact with one from the other prM_3_E_3_. The proteins are re-distributed around the symmetry axes.

## Materials and Methods

### Cells and viruses

Human neuroblastoma SK-N-SH cells (ATCC HTB-11) were maintained in Dulbecco’s Modified Eagle’s Medium with 1000 mg/l glucose (DMEM; Sigma-Aldrich) supplemented with 10 % fetal bovine serum (FBS, Gibco), 0.5 mg/ml penicillin, 500 U/ml streptomycin (penstrep; Lonza Bioscience), 2 mM glutaMAX (Gibco), and non-essential amino acids (NEAA; Gibco). Baby hamster kidney cells (BHK-21, ATCC CCL-10) were maintained in DMEM (Sigma-Aldrich) supplemented with 10 % FBS (Sigma-Alrich). All cells were maintained at +37 °C at a 5% CO2 atmosphere.

TBEV strain Kuutsalo-14 (European subtype; GenBank MG589938.1) was a kind gift from Prof. Olli Vapalahti, University of Helsinki. The virus stock was produced in SK-N-SH cells and titered using a plaque assay as described previously^6^. TBEV strain Hypr (European subtype; GenBank U39292.1) was passaged five times in the brains of suckling mice and once in BHK-21 before its use in the present study. The virus was provided by the Collection of Arboviruses, Biology Centre of the Czech Academy of Sciences (https://arboviruscollection.bcco.cz). TBEV strain Neudoerfl (European subtype; GenBank U27495.1) was passaged several times in the brains of suckling mice, in UKF-NB-4 and BHK-21 cells before its use in the present study. The virus was kindly provided by Prof. Franz Heinz, Medical University of Vienna. The Neudoerfl and Hypr titers were estimated by plaque assay as described previously^35^.

### Production and purification of immature TBEV

For production of Kuutsalo-14 immature particles, SK-N-SH cells were grown to 90 % confluency and, the virus was added in the infection medium (DMEM, 2 % FBS, glutaMAX, penstrep, NEAA, 0.35 uM rapamycin) at a multiplicity of infection of 10. At 22 h.p.i. cells were washed with potassium-buffered saline (PBS) and a fresh infection medium containing 20 mM NH4Cl was added. At 24 h.p.i. the procedure was repeated. The supernatant containing immature TBEV particles was collected at 48 h.p.i. and precleared by centrifugation at 4000 *g* for 5 min. Immature TBEV was pelleted by centrifugation through a 30 % sucrose cushion in HNE buffer (20 mM HEPES pH 8.5, 150 mM NaCl, 1 mM EDTA) at 131,000 x *g* at +4 °C for 2 h. The supernatant was discarded and the pellet was resuspended in HNE, treated with 25 U of benzonase (MerckMillipore) and immediately loaded onto linear glycerol-potassium tartrate gradients (30 % glycerol – 10% glycerol, 35% potassium tartrate (w/v). Following a 2 h centrifugation at 126,500 x *g* at +4 °C, particle-containing light-scattering bands were collected. The samples were concentrated and buffer-exchanged to HNE using Amicon Ultra centrifugal filters (Merck) and irradiated with 25 mJ/cm^2^ of UV_245nm_ to inactivate infectivity. The protein concentration was determined using a Qubit Protein Kit (ThermoFisher), and the protein content was analyzed using sodium dodecyl-sulphate polyacrylamide gel electrophoresis (SDS-PAGE) and immunoblotting as described previously^6^.

For production of Hypr and Neudorfl immature particles, BHK-21 cells were grown to 85 % confluency and infected by TBEV strain Hypr or Neudoerfl at a multiplicity of infection of 1. The cells were incubated in medium (DMEM, 5 % FBS, 25 mM HEPES, pH 7.4) for 18 h (Hypr strain) or 26 h (Neudoerfl strain). The medium was replaced by medium containing NH4Cl (DMEM, 2 % FBS, 25 mM HEPES, 20 mM NH4Cl, pH 7.4) and the cells were incubated for 24 h at +37°C, 5 % CO2. After incubation, the medium was clarified by centrifugation (5,700 x *g*; +4 °C, 20 min) and PEG 8000 dissolved in TNE buffer (20 mM Tris, 120 mM NaCl, 1mM EDTA, pH 8.5) was added to the clarified supernatant. The final concentration of PEG was 8 % (w/v). The particles were then fixed by addition of 0.05 % formaldehyde (v/v, final concentration) and precipitated O/N in an orbital shaker at 130 rpm, +4 °C. The precipitated particles were pelleted at 15,000 x *g*, for 60 min at +4 °C. The pellet was resuspended in 10 ml of TNE buffer containing 8 % PEG 8000 (w/v) and pelleted by centrifugation at 15,000 x *g* for 60 min at +4 °C. The pellet was resuspended in 3 ml of TNE buffer, RNAse A was added (10 µg/ml, final concentration), and incubated for 15 min at 15 °C. The suspension was centrifuged at 15,000 g, 10 min, +4 °C, and the supernatant was loaded on 10-35% (w/v) potassium tartrate step gradient and centrifuged at 175,600 x *g*, 2 h, +4 °C. The light-scattering band was collected. The sample was buffer exchanged into TNE buffer by serial dilution and concentration using Amicon Ultra centrifugal filters (Merck).

### In vitro maturation

Each 15 µL in vitro maturation reaction contained 5 µg of purified, UV-inactivated immature Kuutsalo-14 TBEV in HNE adjusted to the indicated pH using 1 M 2-(N-morpholino)-ethanesulfonic acid hydrate (MES, pH 5.0). Reactions were supplied with 30 mM CaCl2 and 2U human recombinant furin (Thermo-Fisher, RP-062) and incubated at +30 °C for 16 h. After incubation, half of each reaction was mixed with 4x Laemmli sample buffer and proteins were resolved in 4-20% gradient SDS-PAGE. Protein bands were visualised using stain-free imaging using Biorad GelDoc EZ, and specific prM and M bands were visualised using immunoblotting with an anti-M antibody as described previously^6^.

### CryoEM sample preparation

The samples were vitrified in liquid ethane on glow-discharged 200 copper mesh R1.2/1.3 Quantifoil holey carbon coated grids with 2 nm continuous carbon on top (Jena Bioscience) using a Leica EM GP plunger at 85 % humidity with 1.5 s blotting time. Both TBEV strain Neudoerfl and Hypr samples were vitrified in liquid ethane on holey carbon coated copper grids (Quantifoil 2/1, mesh 300, Quantifoil Micro Tools GmbH) using Vitrobot Mark IV (Thermo Fisher). Samples were stored under liquid nitrogen until use.

### CryoEM data collection

Kuutsalo-14 data were collected at the SciLifeLab CryoEM Infrastructure Unit (Solna, Sweden) using FEI Titan Krios microscope (Thermo Fisher) operating at 300 kV equipped with a Gatan K3 detector in counting mode at nominal magnification 81,000 x resulting in a sampling rate of 1.075 Å/px. Movies were recorded in counting mode at a total dose of 43 e^-^/Å^2^ distributed over 50 frames using defocus range −0.7 to −2.7 um (step 0.2 um). EPU software (Thermo Fisher) was used for data acquisition. In total, 69964 movies were collected. TBEV strain Hypr was collected on a Titan Krios microscope (Thermo Fisher) operating at 300 kV aligned for parallel illumination in nanoprobe mode equipped with a Falcon3EC direct electron detector. The micrographs were collected in integration mode at a nominal magnification of 75,000 x resulting in a 1.08 Å pixel size on the detector. The defocus range applied during the acquisition was −3 to −1 µm and the total dose during the 1 s acquisition was 69 e^-^/Å^2^. The dose fractionated acquisitions were saved as 39 fraction movies. EPU software was used for the data acquisition. In total 2,262 movies were collected. TBEV strain Neudoerfl was collected on a Titan Krios microscope operating at 300 kV aligned for fringe-free imaging and equipped with a Gatan K3 direct electron detector behind an energy filter (BioQuantum K3, Ametek) with 10 eV slit inserted. The micrographs were collected in counting mode at a nominal magnification of 105,000 x resulting in a 0.8336 Å pixel size on the detector. The nominal defocus range applied during the acquisition was −3 to −1 µm and the total dose during the 2 s acquisition was 40 e^-^/Å^2^. The dose fractionated acquisitions were saved as 40 fraction movies. SerialEM software^36^. was used for the data acquisition utilising the beam-tilt compensation upon image shift. In total 11,246 movies were collected. Data for both Hypr and Neudoerfl strains were collected at Cryo-electron microscopy and tomography core facility CEITEC MU (Brno, Czech Republic). See Table 1 for full data collection statistics.

### Image processing

Kuutsalo-14 image processing was performed at the Finnish Center for Scientific Computing (CSC) supercomputing cluster using Scipion 3.0 framework and Cryosparc^21,24^. Movies were aligned using MotionCor2 implemented in Relion 3.1^37,38^, and movies with per-frame motion exceeding 5 pixels were rejected using Xmipp movie maxshift^39^. Contrast transfer functions were calculated using gctf and CTFFind4, and micrographs with resolution discrepancies beyond 3.0 Å, as well as with low-resolution (worse than 6 Å), astigmatism (> 500 Å) were rejected using CTF consensus^39–41^. Particles were picked and extracted with a box size of 900 px using Xmipp3 and several rounds of 2D-classification were done using Relion 3.1.2. The selected particles (96,433) were used for initial model generation using stochastic gradient descent method as implemented in Relion 3.1.2^37^. Particles were subsequently 3D classified and refined in Relion 3.1 or in Cryosparc using a 650 Å spherical mask to a final resolution of 7.10 Å according to the FSC_0.143_ criterion. For localized reconstruction of prM_3_E_3_ spikes, sub-particles were defined and extracted using localised reconstruction package integrated into Scipion^42^ and exported using Relion export particles function. The sub-particles were imported into Cryosparc v4.2.1 and were reconstructed using iterative *ab initio* model generation, 3D classification, and refinement using a soft segment mask that included a single spike with the membrane part. The final resolution of the reconstructed spike was 3.89 Å according to the FSC_0.143_ criterion. The local resolution of prM_3_E_3_ map was estimated using Cryosparc and the map was locally sharpened using Xmipp3 or minimally sharpened using Relion 3.1 post-processing function. Both the locally and the minimally sharpened maps were used for model building.

The TBEV strain Hypr data was processed as follows. The collected dose-fractionated movies were aligned using MotionCor2^38^ and the sums of dose-weighted fractions were saved as individual micrographs. CTF estimation was performed using Gctf v1.06^40^. The initial set of particles was manually picked on non-binned micrographs and the crYOLO neural network was trained on this subset. Automatic particle picking was performed on micrographs using crYOLO^43^. For initial 2D classification, the particles were extracted and down-sampled from 768 px box size to 128 px (final pixel size 6.48 Å). Several rounds of 2D classification were performed in Relion 4.0.0^44^. The selected 18,160 particles were re-extracted and down-sampled to box size of 512 px (pixel size 1.62 Å). The stochastic gradient descent method as implemented in Relion 3.1.2^37^ was used for initial model generation. Refinement using the initial model lowpass filtered to 40 Å was done in Relion v3.1.2 with icosahedral symmetry applied, using only a spherical mask of diameter 650 Å. No further 3D classification or masked refinement improved the map resolution or quality.

The final map was masked by a threshold mask and B-factor sharpened in the postprocessing procedure using Relion 3.1.2. The final resolution was estimated using the FSC_0.143_ criterion as 8.64 Å.

The TBEV strain Neudoerfl data was processed as follows. The collected dose-fractionated movies were aligned using MotionCor2 and the sum of dose-weighted fractions were saved as individual micrographs. CTF estimation was performed using Gctf v1.06. The initial set of particles was hand-picked on 10 x binned micrographs and crYOLO neural network was trained on this subset. Automatic particle picking was performed on 10 x binned micrographs using crYOLO and the resulting coordinates were corrected by the binning factor to match the particle positions on the unbinned micrographs. For initial 2D classification the particles were extracted and down-sampled form 960 px box-size to 128 px (final pixel size 6.25 Å). Several rounds of 2D classification were performed in Relion 4.0.0 to exclude false positive particles picked by crYOLO. The resulting 36,236 particles were reextracted and down-sampled to box-size of 512 px (pixel size 1.56 Å). Refinement using lowpass filtered (40 Å) initial model from a previous refinement of TBEV Hypr was done in Relion v 3.1.2 with icosahedral symmetry applied, using only a spherical mask of diameter 650 Å. No further 3D classification or masked refinement improved the map resolution or quality. The final map was masked by a threshold mask and B-factor sharpened in the Relion 3.1.2 postprocessing procedure. The final resolution was estimated using the FSC_0.143_ criterion as 7.15 Å. To improve the resolution of the spike-trimers, single spike-trimers were extracted in 300 px box from the original micrographs by a modified version of localised reconstruction^23^. In total 1,639,578 sub-particles were extracted and subjected to initial 3D refinement using local searches around the already known orientations. A soft segment mask that included a single spike-trimer together with the membrane was applied in the refinement step. After refinement 3 rounds of 3D classification were performed, dividing the particles into 40 classes. The orientational search was omitted during the classification and the orientations from the previous refinement step were used. The selected classes included 552,993 particles, which were subjected to 3D refinement. This was followed by anisotropic magnification estimation, 3^rd^ and 4^th^ order aberration estimation and defocus refinement per particle in Relion 3.1.2. Finally, the volumes were reconstructed using relion_reconstruct with applied Ewald sphere correction. The final map was masked and B-factor sharpened. The final resolution was estimated using the FSC_0.143_ criterion as 4.03 Å.

### Model Building

The homology model of Kuutsalo-14 prM_3_E_3_ was generated using I-TASSER^45^ with BinJV prM_3_E_3_^19^ (PDB ID: 7l30) as a template. Membrane-associated domains of prM (residues 112-161) were replaced with membrane-associated domains of M from mature TBEV^6^ (PDB ID: 7z51). The resultant model was flexibly fitted into the prM_3_E_3_ maps using ISOLDE integrated into ChimeraX^46,47^. The model was real-space refined using Phenix version 1.20^48^, after which clashes were fixed in ISOLDE with model-based distance and torsions constraints imposed. For TBEV strain Neudoerfl an Alphafold v2.2^49^ generated model of the E-prM complex multimer was manually rigid body fitted into the post-processed electrostatic-potential map of the prM_3_E_3_ spike using UCSF Chimera^50^. The ectodomain of the E-protein and the prM fitted well and manual adjustments were done in Coot^51^, followed by real-space refinement in Phenix v1.20. The membrane part of the E-protein was modeled using BinJV prM_3_E_3_^19^ (PDB ID: 7l30) E-protein transmembrane domain as template and the membrane part of the prM was modelled using TBEV M protein^5^ (PDB ID: 5O6A) as a template. Because of the low resolution of the map in the membrane region, the sidechains of the amino acid residues in this region were stripped, and secondary structure restraints were applied before Phenix real-space refinement. The final model containing the joined ectodomains and transmembrane domains was iteratively refined using Phenix real-space refinement and the refined model was manually inspected and corrected in Coot, till final convergence. The geometry of all models was continuously monitored using MolProbity^52^.

## Acknowledgments

We thank Dr. Pasi Laurinmäki of the Instruct-ERIC Centre Finland and the Biocenter Finland National CryoEM Facility, the CryoEM Swedish National Facility funded by the Knut and Alice Wallenberg, Family Erling Persson and Kempe Foundations, SciLifeLab, Stockholm University and Umeå University, and the CSC-IT Center for Science Ltd. for providing reagents, technical assistance, and facilities to carry out the work. We thank Prof. Olli Vapalahti for the kind gift of the TBEV Kuutsalo-14 isolate and Saana Haarma, Dr. Suvi Kuivanen, Irina Suomalainen and Sanna Mäki for excellent technical help. We thank Prof. Franz Heinz for the kind gift of TBEV strain Neudoerfl.

The research was funded by the Swedish Research Council, reference number 2018-05851 (to SJB); the Academy of Finland, grant number 315950 (to SJB); the Sigrid Juselius Foundation, grant number 95-7202-38 (to SJB); the Jane and Aatos Erkko Foundation (to SJB); European Union’s Horizon 2020 Research and Innovation Programme under the Marie Skłodowska-Curie grant agreements number 799929 (to MA); University of Helsinki Research Foundation (to LIAP); University of Helsinki Doctoral School in Integrative Life Science (to LIAP); Helsinki Institute of Life Sciences (to SJB), the project “National Institute of Virology and Bacteriology” (Programme EXCELES, ID Project No. LX22NPO5103) - Funded by the European Union - Next Generation EU (to PP and DR) and the Czech Science Foundation Grant GX 19-25982X (to PP). We acknowledge Cryo-electron microscopy and tomography core facility CEITEC MU of CIISB, Instruct-CZ Centre, supported by MEYS CR (LM2023042) and European Regional Development Fund-Project „UP CIISB“ (No. CZ.02.1.01/0.0/0.0/18_046/0015974).

## Author contributions

M.A., S.J.B., and P.P. designed research; M.A., L.I.A.P., P.F.-P., P.S., D.R., J.N, L.S. and T.F. performed research; M.A., A.D. S.J.B., T.F. and P.P analysed data; M.A. drafted the paper and all authors contributed to finalising the paper.

## Competing Interests Statement

The authors declare no competing financial interests.

## Notes

### Competing Interest Statement

The authors have declared no competing interest.

