## Supplementary material for "The structure of immature tick-borne encephalitis virus": Figures S1-S8

### Supplementary Information

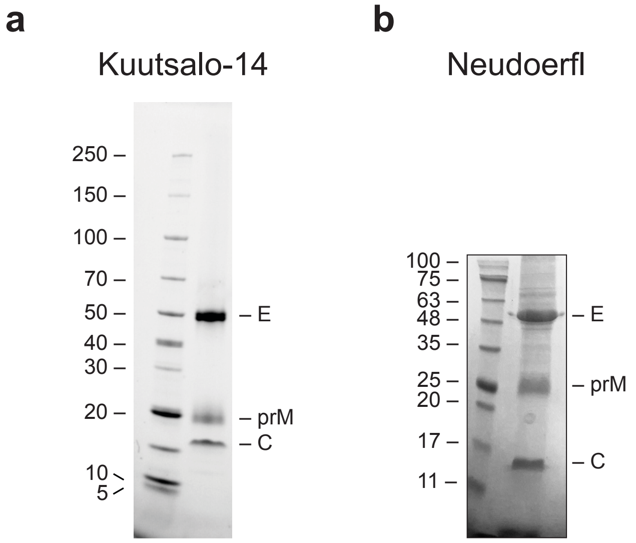


*Figure S1.* ***Purity of immature TBEV preparations.*** *SDS-PAGE of purified immature particles shows the 3 major protein components, E, prM, and C for Kuutsalo-14 (**a**) and Neudoerfl (**b**). Molecular weight markers are shown on the left.*

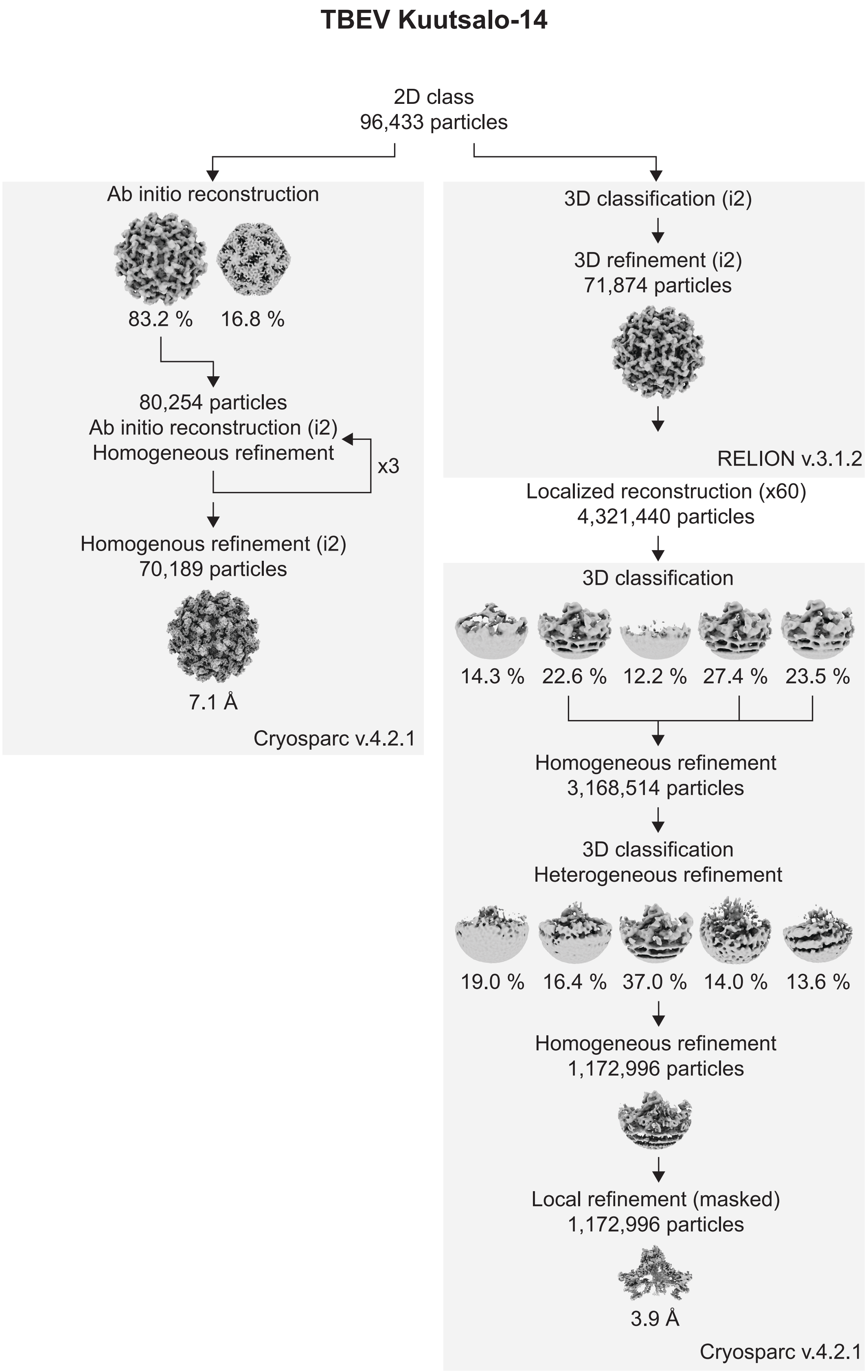


*Figure S2.* ***CryoEM data processing flow chart of immature Kuutsalo-14 TBEV.*** *Flowchart of processing steps of immature Kuutsalo-14 going from selected 2D class to icosahedral reconstruction of immature particles, and to localised asymmetric reconstruction and refinement of trimeric* $prM_3E_3$ *spike.*

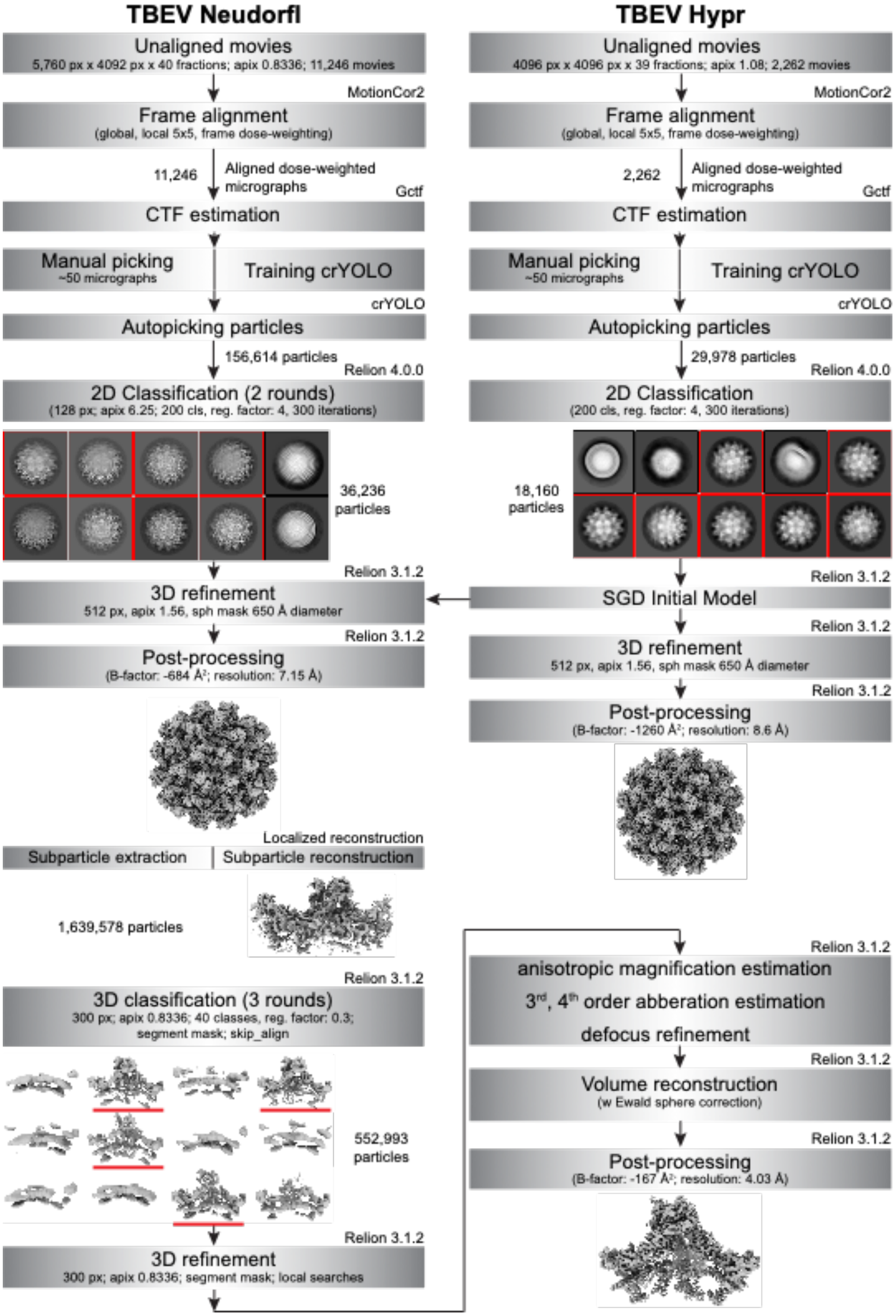


*Figure S3.* ***CryoEM data processing flow charts of immature Neudoerfl and Hypr TBEV.*** *Flowcharts of processing steps of immature Neudoerfl and Hypr TBEV starting from image pre-processing, particle picking, and classification, to the icosahedral reconstructions of immature particles for both strains and to localised asymmetric reconstruction and refinement of trimeric prM$_3$E$_3$ spike of the Neudoerfl strain.*

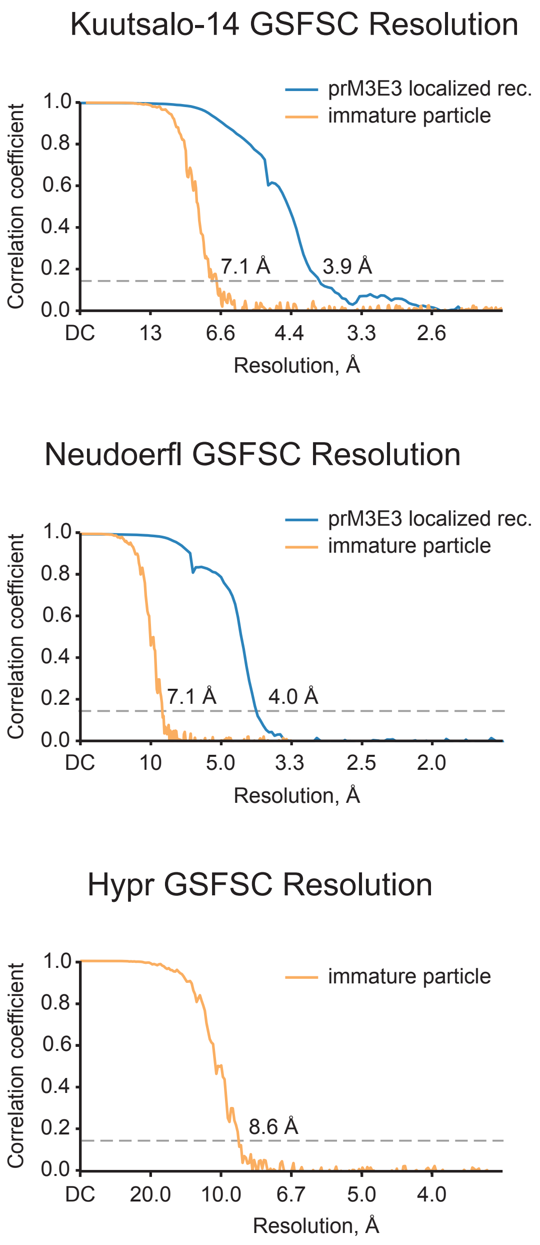


*Figure S4.* ***Fourier Shell Correlation curves of the half-reconstructions using gold-standard refinement in RELION and CryoSPARC.*** *The grey dashed line indicates a 0.143 FSC cutoff and approximate resolutions at this cutoff are indicated. The X-axis extends to the Nyquist resolution of the collected dataset. FSC curves are shown for masked half-maps.*

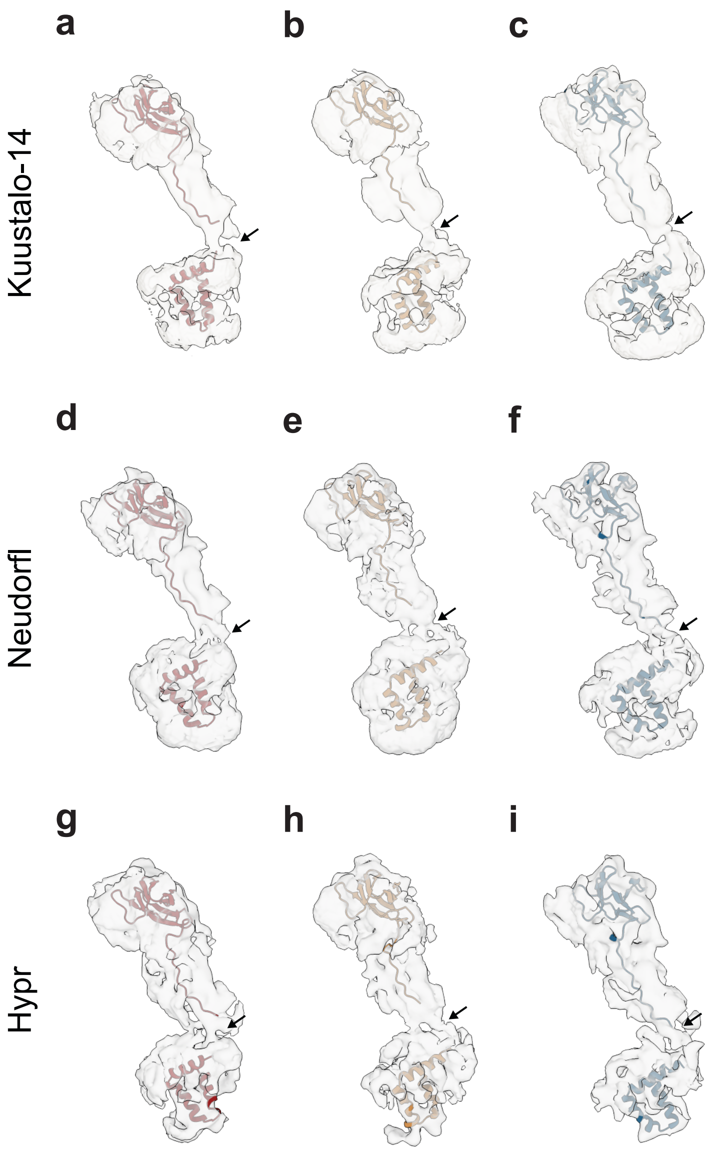


*Figure S5.* ***prM linker density observed in icosahedral reconstructions of immature TBEV.*** *Sections of whole particle density maps surrounding rigidly fitted atomic models of Kuutsalo-14 (**a-c)**, Neudoerfl (**d-f**), and Hypr (**g-i**). Individual prM chains are shown for each prM copy within an asymmetric unit with linker densities indicated with arrows.*

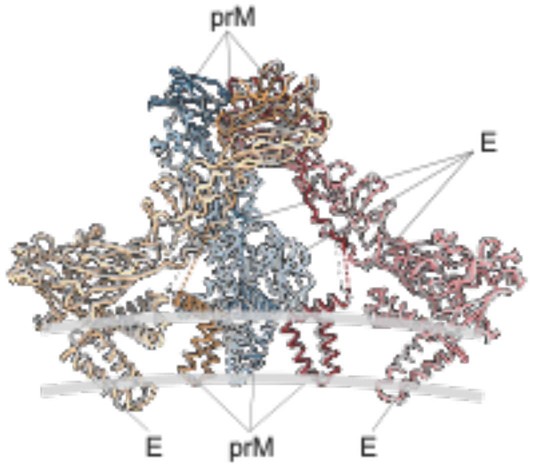


*Figure S6.* ***Comparison of $prM_3E_3$ models of Kuutsalo-14 and Neudoerfl.*** *An overlay of Kuutsalo-14 (colouring convention as in Fig. 3) and Neudoerfl (grey) $prM_3E_3$ models in string representation. E and prM proteins are indicated, and grey lines mark the position of the lipid bilayer.*

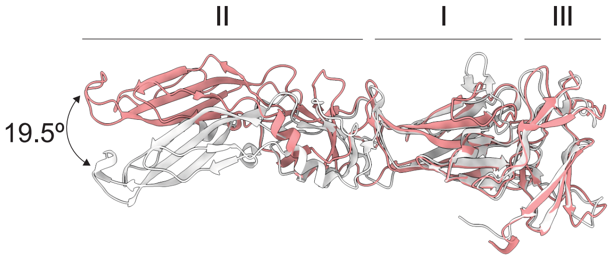


*Figure S7.* ***Comparison of sE of cryoEM and X-ray prME models.*** *An overlay of E protein ectodomain from Kuutsalo-14 prM$_3$E$_3$ (red) and from Neudoerfl (pr/sE)$_2$ (grey; PDB ID: 7QRE). Domains I, II, and III are indicated, and a 19.5° difference in the positions of domains II is indicated.*

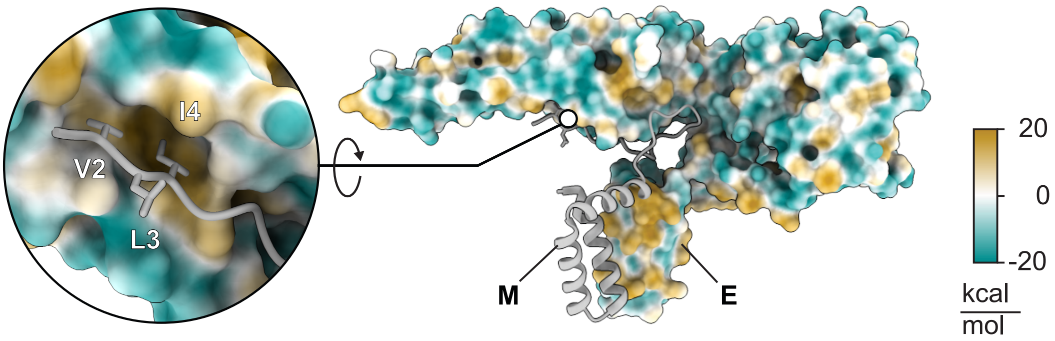


*Figure S8.* ***A hydrophobic zipper downstream of the furin cleavage site is maintained in the E-M dimer of the TBEV virion.*** *A surface representation of E coloured by hydrophilicity and a cartoon representation of M from TBEV virion (PDB ID: 7z51) show a stretch of hydrophobic residues Val2-Leu3-Ile4 on M docked into a hydrophobic pocket of E, proximal to the membrane. Residues Val2, Leu3, and Ile4 in M correspond to residues Val90, Leu91, and Ile92 of prM involved in stabilisation of the furin cleavage site in the immature TBEV.*
